# Reversible inhibition of medial septum interneurons abolishes initial acquisition of new place-reward associations and reveals a progressive reduction in dependence on medial septal function

**DOI:** 10.64898/2026.09.25.754563

**Authors:** Zahra Alizadeh, Janina Ferbinteanu, Michael Eckert, Justin Shobe, Robert J. McDonald, Bruce L. McNaughton

## Abstract

Hippocampal place cells encode spatial information within a theta-organized temporal framework paced by the medial septum (MS). Lesions or inactivation of the MS impair spatial learning and disrupt hippocampal theta rhythmicity, long-term potentiation, and plasticity-related gene expression, while often sparing spatially selective firing of CA1 neurons (but not CA3 neurons). Although many rodent studies argue against classical systems consolidation by showing persistent hippocampal dependence of spatial memories, other work has suggested that training structure, spaced vs. massed trials, can influence this dependence.

To investigate this possibility, we used DREADD-based, reversible, inhibition of MS GABAergic neurons, some of which are known to project to hippocampus. DREADD activation reduced hippocampal theta power and spatial information encoded by hippocampal neurons while largely preserving overall neuronal firing. We then examined the effect of inhibition of medial septal GABAergic neurons on learning the correct goal arm in an 8-arm maze when the correct arm was randomly changed each day. On each day, the mice were given two ‘forced-choice’ trials indicating the correct arm and then 5 free-choice trials (without correction) and were always started from an opaque cylinder in the maze center, in a random orientation, to eliminate possible use of a simple response strategy. When the correct goal arm varied randomly daily, MS cell inhibition severely and repeatedly impaired acquisition. In contrast, when the correct goal arm was subsequently fixed across days, performance was initially impaired but became resistant to DREADD activation over time. Once performance reached asymptote, the goal arm was switched to a new (random) position. In this case, learning was no longer impaired by DREADD activation. This generalization suggests that repeated fixed-goal training in the same environment induces a broader reorganization of the neural systems supporting spatial behaviour. One possibility is that mice develop a septal-independent task-solving strategy that supports both retrieval of established goal locations and acquisition of new ones within a stable environment. This transition could reflect the gradual adoption of a ‘non-spatial’ strategy such as visual pattern recognition, or the use of a directional cue. Moving the most salient distal cue in the environment (a single room light) produced a substantial change in the visual configurations seen from the arm entry points, and disrupted performance; however, it did not produce a consistent directional shift relative to the cue, suggesting that the light contributed to an integrated visuo-spatial representation rather than serving as the reference frame origin. We conclude that the mice switched from a hippocampus/MS-dependent spatial strategy to a non-hippocampus-dependent visual pattern matching solution.

## Introduction

The hippocampus (HPC) plays a central role in spatial learning through the activity of place cells, neurons that fire selectively at specific locations in the environment (O’Keefe and Dostrovsky, 1971) and are modulated by sensory feature changes or events at a given location (rate remapping). This spatially tuned activity is temporally organized by the hippocampal theta rhythm, such that, as animals move through space, place cells exhibit theta modulation and phase precession, shifting their spike timing systematically across the theta cycle (O’Keefe & Recce, 1993). The generation and maintenance of hippocampal theta depend on the medial septum (MS), which has long been proposed to act as a pacemaker of hippocampal rhythmicity (Buzsáki et al., 1983; Freund and Antal, 1988; Köhler et al., 1984; Stewart and Fox, 1990; Tóth et al., 1993; Wainer et al., 1985; Wang, 2002). Early electrophysiological studies demonstrated that the hippocampus exhibits prominent rhythmic activity during arousal and active behaviour (Green and Arduini, 1954), establishing theta as a key feature of hippocampal network dynamics. Subsequent lesion studies showed that disruption of medial septal input abolishes hippocampal theta oscillations and impairs spatial memory (Allen and Crawford, 1984; Bland et al., 1996; Givens and Olton, 1990, 1994; Winson, 1978), highlighting the importance of MS for learning and memory. Further work demonstrated that the MS coordinates hippocampal activity via GABAergic and cholinergic inputs that regulate interneurons and pyramidal cell firing (Bland and Colom, 1993; Smythe et al., 1992).

In addition to its role in network dynamics, septal input has been strongly linked to synaptic plasticity mechanisms underlying memory formation. Disruption of septal activity impairs long-term potentiation (LTP) and plasticity-related processes in the hippocampus (Hölscher et al., 1997; Kubik et al., 2007; M’Harzi et al., 1987), possibly by disrupting theta oscillations (Buzsáki, 2002; Leung, 1998) or disruption of cholinergic input, which has been shown to modulate synaptic transmission and plasticity within hippocampal circuits (Dannenberg et al., 2017; Hasselmo and Schnell, 1994). Consistent with this view, more recent work has shown that disrupting spike timing at the theta timescale impairs short-term spatial working memory, while sparing spatial reference memory and hippocampal replay, further emphasizing the importance of precise temporal coordination for some aspects of memory (Joshi et al., 2026).

Despite these deficits, hippocampal firing rates and spatially selective activity, particularly in CA1, can remain relatively preserved following septal disruption. Reversible inactivation studies demonstrated that place-specific firing persists in CA1 but not CA3 even when theta is reduced and behavioural performance is impaired (Mizumori et al., 1989; Mizumori et al., 1990; Mizumori et al., 1992), suggesting that theta is not required for the expression of spatial representations *per se*, but rather for their temporal organization and stability. Consistent with this dissociation, selective disruption of the temporal organization of hippocampal place-cell activity within theta cycles impairs theta sequences and predictive spatial representations while preserving single-cell spatial tuning and rate coding, resulting in impaired flexible memory-guided navigation (Liu et al., 2023).This dissociation has been further supported by work showing that reduction of theta disrupts the spatial periodicity of grid cells, theta phase precession, spike timing coordination, and the temporal organization of hippocampal firing, without abolishing spatial selectivity (Brandon et al., 2011; Hinman et al., 2011; Koenig et al., 2011; Robinson et al., 2024; Wang et al., 2015). Furthermore, pharmacological disruption of medial septal circuits demonstrates that the effects may depend on the method used. Complete septal inactivation with tetracaine or muscimol disrupts hippocampal theta and reduces the spatial firing of place cells, whereas gabazine abolishes theta while largely preserving location-specific place-cell firing. Despite this preservation, gabazine still impairs spatial navigation, indicating that disruption of septo-hippocampal temporal coordination alone is sufficient to impair behaviour (Bolding et al., 2020). Together, these findings support a model in which septal input does not generate spatial representations directly, but instead organizes the timing, structure, and informational content of hippocampal coding through both rhythmic coordination and cholinergic modulation of synaptic plasticity (Buzsáki, 2002).

At the systems level, the role of the hippocampus in memory has been classically framed by theories of systems consolidation, which propose that newly acquired memories are initially hippocampus-dependent but become gradually reorganized into distributed cortical networks over time (Frankland and Bontempi, 2005; Squire & Zola-Morgan, 1991). Supporting this framework, studies using brain activity mapping and cortical inactivation (anterior cingulate cortex) have demonstrated a time-dependent shift in neural activity from the hippocampus to neocortical regions during memory retrieval (Bontempi et al., 1999; Frankland et al., 2004). However, a substantial body of rodent work has challenged this view by demonstrating persistent hippocampal dependence of spatial memories, even at remote time points. Lesion studies established that the hippocampus is critical for spatial learning (Morris et al., 1982; Sutherland et al., 1982), and subsequent work showed that disruption of hippocampal output pathways similarly impairs spatial memory (Sutherland and Rodriguez, 1989). More recent studies have reported that hippocampal lesions impair both recent and remote spatial memory, suggesting that retrieval may continue to rely on hippocampal circuitry regardless of memory age (Sutherland et al., 2001; Martin et al., 2005; Clark et al., 2005). Consistent with this view, critical reviews have argued that evidence for temporally graded retrograde amnesia in spatial tasks is weak or inconsistent, and that the hippocampus may remain necessary for detailed spatial representations (Sutherland et al., 2010; Eichenbaum, 2000). These findings raise the possibility that, at least for spatial tasks, hippocampal involvement may remain necessary for memory expression, calling into question a strictly time-dependent transfer of memory dependence to cortical structures.

In addition, other work has emphasized that the degree of hippocampal dependence may be strongly influenced by training structure, rather than time alone. In a contextual fear conditioning study, hippocampal lesions abolished performance after massed training (12 shocks in a single session), whereas the behaviour was preserved after lesions in animals given the same number of shocks distributed over 12 days (Sutherland and Lehmann, 2011; Sutherland et al., 2010). It was suggested that the distributed training protocol provided more opportunities for post-trial memory reactivation, which may have enabled the development of a hippocampus-independent representation. Such representations may be supported by cortical circuits and/or other parallel learning and memory systems that acquire information more gradually than the hippocampal system. More broadly, the concept of distributed learning systems proposes that memory is supported by parallel and interacting networks, whose relative contributions depend on experience and training conditions (Sutherland et al., 2010; White and McDonald, 2002; McDonald et al., 2017). Under this framework, repeated exposure allows cortical circuits to extract regularities and form stable representations while also allowing other parallel learning and memory networks to provide alternative solutions.

Despite extensive evidence that medial septal input is critical for spatial learning and the temporal organization of neural activity, it remains unclear whether this dependence is fixed or dynamically modulated by experience. In particular, while disruption of septo-hippocampal interactions reliably impairs learning under conditions requiring flexible spatial encoding, it is unknown whether repeated training can reduce reliance on medial septal function and allow behaviour to be supported by alternative, possibly neocortical, neural systems.

We first verified the physiological consequences of medial septal inhibition using silicon-probe recordings, confirming that the manipulation reduced hippocampal theta power and spatial information while largely preserving neuronal firing. We examined how the role of medial septal interneurons in spatial learning changes with different types of training. Using DREADD-based inhibition of MS interneurons in an 8-arm maze, we compared a condition in which the rewarded arm changed every day with one in which it stayed the same across days.

## Results

### Theta power and spatial information of hippocampal place cells were reduced after DCZ administration while average firing rate and multi-unit activity remained stable

To assess the effect of inhibition of medial septal inhibitory interneurons on the rhythmic organization of hippocampal activity, we examined MS DREADD (pAAV-hDlx-GiDREADD-dTomato-Fishell-5) induced changes in oscillatory dynamics, multi-unit activity (MUA), and spatial coding of pyramidal cells in both CA1 and in the overlying neocortex, which exhibits the development of spatial codes (Mao et al.,2017; Esteves et al., 2021; Navratilova et al., 2026) that depend initially on hippocampus but become independent of it over time (Esteves et al., 2021). Two head-restrained mice were trained to walk on a rotating drum that updated a visual virtual reality (VR) display (Figure 1A, B, D). Recordings were made with silicon probes spanning the dorsal cortex and dorsal and ventral hippocampus.

**Figure 1.**
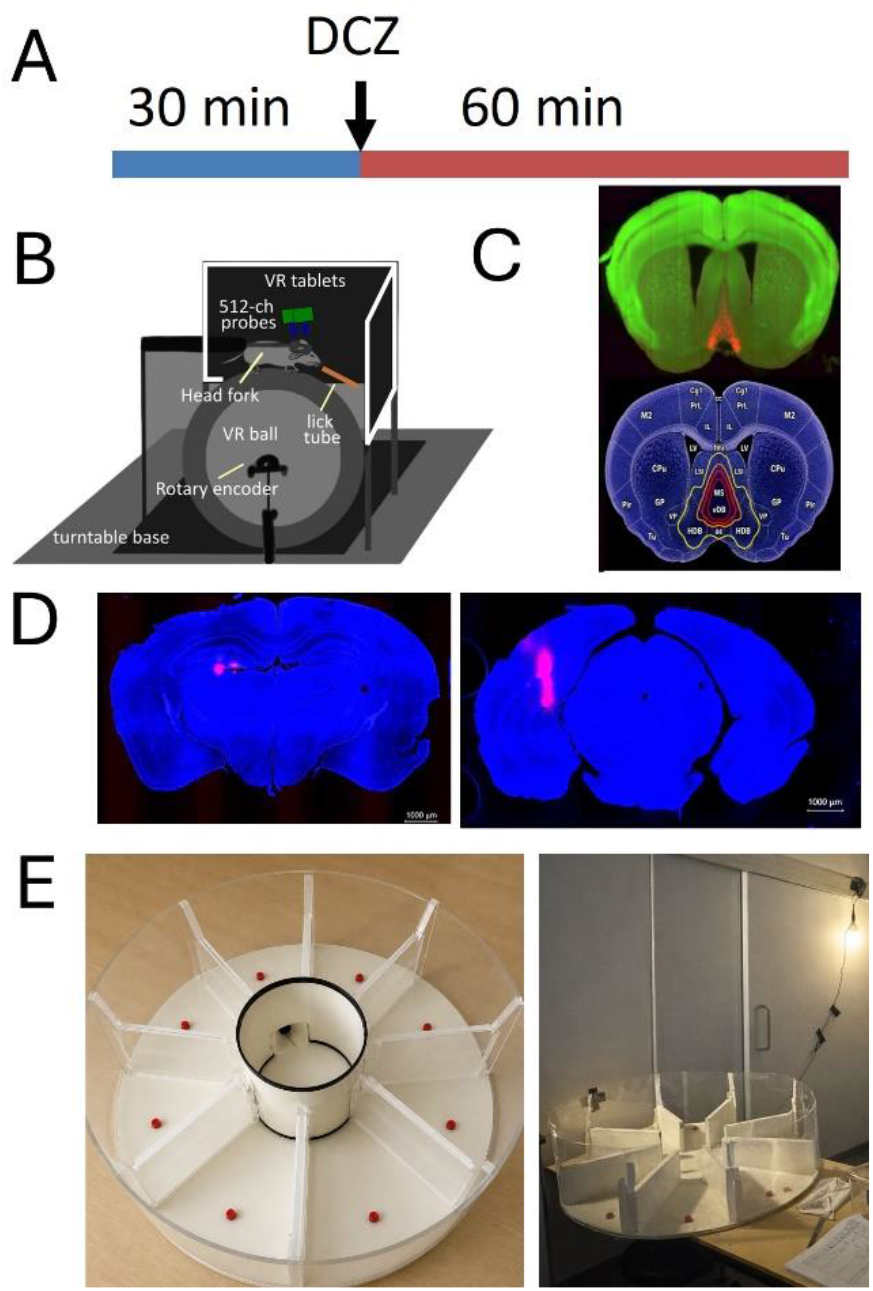
Experimental design, behavioral and electrophysiological apparatus, and histological verification. (A) Schematic of the electrophysiological recording paradigm. Animals ran on the virtual reality treadmill during a 30 min baseline recording epoch, followed by DCZ administration (0.5 mg/kg) and an additional 60 min recording period during locomotion. (B) Schematic of the virtual reality treadmill apparatus used for electrophysiological recordings. Head-fixed mice ran on a rotating drum while navigating a virtual reality environment displayed on surrounding monitors. The setup included a head-fixation bar, rotary encoder, lick tube for reward delivery, and surrounding visual displays. (C) Top, representative expression pattern of medial septum DREADD for the behavioral experiments (green, tissue autofluorescence; red, viral expression). Bottom, summary of expression from MS injections in 22 mice superimposed on the median atlas section (contour lines: yellow, 25%; orange, 50%; red, 75%; maroon, 90%). Medial septal input to the hippocampus comprises three major projections: GABAergic, cholinergic, and glutamatergic. GABAergic septal neurons are the principal pacemakers of hippocampal theta and coordinate rhythmic timing across hippocampal circuits. Cholinergic septal neurons regulate excitability, plasticity, and attention-related modulation of hippocampal processing. Glutamatergic septal neurons provide excitatory drive (Müller and Remy, 2018) (D) Representative coronal sections showing DREADD expression and silicon probe placement for electrophysiological recordings (blue, DAPI; red, viral expression and probe track). (E) Behavioral apparatus and testing environment for the radial arm maze task. Left, overhead view of the 8-arm radial maze with a central opaque start cylinder and reward positioned at the distal end of each arm. Right, experimental room showing the maze setup and the dominant distal light cue used for cue-shift manipulations.

Spectrogram analysis of hippocampal local field potentials revealed a marked reduction in theta-band power following DCZ administration (0.5 mg/kg S.C.; Figure 2A-C). Despite this reduction in theta oscillations, overall neuronal firing rates remained largely preserved following DCZ administration. The MUA power spectrum showed a reduction in theta-range power after DCZ, without a broad suppression of power across frequencies (Figure 2D). Autocorrelograms of representative CA1 pyramidal neurons showed reduced theta-periodic modulation following DCZ administration (Figure 2E–F).

**Figure 2.**
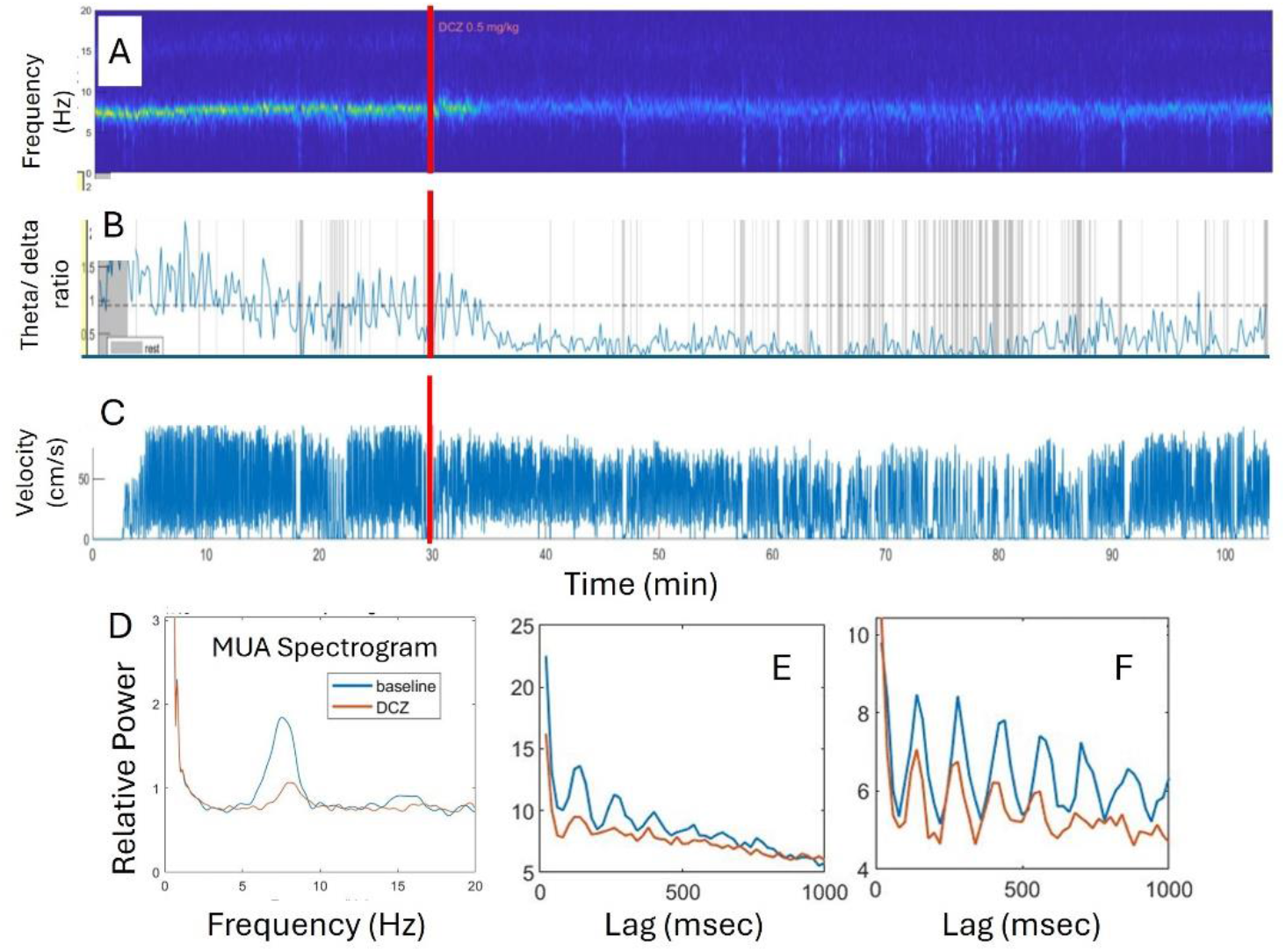
Effects of DCZ administration on hippocampal theta activity and neuronal firing properties. (A) Representative hippocampal local field potential (LFP) spectrogram showing theta power (6–10 Hz) across time. The red vertical line indicates the time of DCZ administration (0.5 mg/kg). (B) Corresponding theta/delta power ratio (6–10 Hz / 1–4 Hz) across the recording session, aligned to the spectrogram in (A). (C) Running velocity across the same recording period. (D) Multi-unit activity (MUA) power spectrum during baseline (blue) and after DCZ administration (orange). (E-F) Autocorrelograms of two representative CA1 pyramidal neurons during baseline (blue) and after DCZ administration (orange).

### Single neuron firing rate was preserved while spatial information was reduced following DCZ administration

To examine the impact of MS inhibition on hippocampal spatial coding, we analyzed single-unit firing rates and spatial information content in both dorsal and ventral CA1 and overlying neocortex (parietal and visual cortex). The mean firing rates of neurons, including both pyramidal cells and interneurons, did not change significantly following DCZ administration. Although some variability was observed at the level of individual units, comparisons between pre- and post-DCZ conditions revealed no consistent or systematic reduction in firing rate in either the dorsal or ventral hippocampus (Figure 3A-B and Supplementary Figure 1C-D).

**Figure 3.**
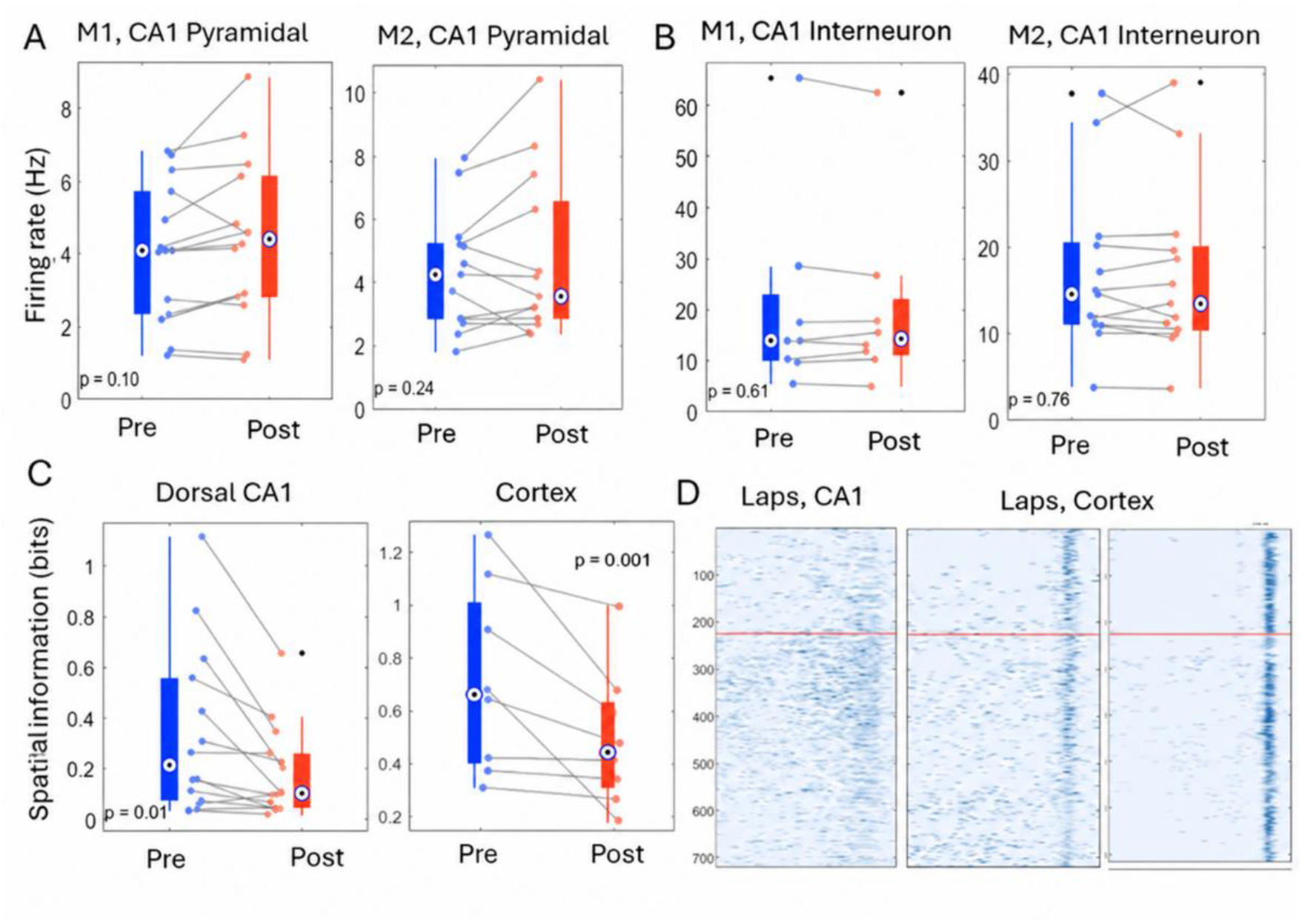
Effects of medial septum inhibition on firing rate and spatial coding in hippocampus and cortex. (A) Firing rates of dorsal CA1 pyramidal cells recorded before (Pre, blue) and after (Post, red) DCZ administration in two representative mice (Mouse1 and Mouse 2). Each point represents one unit, with paired lines connecting pre- and post-DCZ values for the same cell. Boxplots show the distribution, median, and mean firing rate across units. (B) Firing rates of dorsal CA1 interneurons recorded before (Pre, blue) and after (Post, red) DCZ administration in the same two mice. (C) Spatial information of dorsal CA1 pyramidal neurons (left) and cortical neurons (right) before (blue) and after (red) DCZ administration. (D) Representative lap-by-lap firing maps of dorsal CA1 (left) and cortical (right) neurons before and after DCZ administration. Each row represents a single lap, and firing activity is plotted as a function of position along the track. The horizontal red line indicates the time of DCZ administration.

In contrast, spatial coding was significantly affected. Spatial information content of CA1 pyramidal neurons decreased following DCZ administration in the dorsal and ventral hippocampus. While the magnitude of this effect varied across subjects, a consistent reduction in spatial information was observed, indicating that neuronal firing became less informative about the animal’s position in VR (Figure 3C-D and Supplementary Figure 1E). A similar effect was observed in neocortex (Figure 3C-D). Together, these results indicate that inhibition of MS interneurons reduces the quality of hippocampal spatial representations while leaving overall neuronal firing largely preserved across both pyramidal cells and interneurons. This dissociation suggests that septal input is necessary to maintain the structure and informational content of hippocampal activity, rather than simply regulating firing rates. The reduced spatial information in cortex is presumably an indirect effect of the hippocampal disruption, since spatially selective cortical firing in a novel environment depends on an intact hippocampus (Mao et al.,2017; Esteves et al., 2021).

### Medial septum inhibition impaired spatial memory performance

To assess the effect of disrupting the temporal organization of MS interneurons activity on spatial memory, MS DREADD-expressing mice were trained on an 8-arm radial maze in which the rewarded arm was randomly assigned each day. Reward arms were counterbalanced across mice on each day. Two cohorts of MS-DREADD transfected mice were trained on this paradigm for approximately 40 days, during which time performance improved steadily to well above chance or reached asymptote: Cohort 1 (n = 6 males, n = 6 females) and Cohort 2 (16 males). An additional cohort of 4 mice without DREADD transfection received DCZ to control for nonspecific effects. Following completion of training, mice entered the testing phase, DCZ was administered orally in gelatin pellets (Ferrari et al., 2022) on alternating days. Gelatin without DCZ was administered on the intervening control days. Except for the final experiments (Figure 6 C), mice were given two forced-choice ‘information’ trials on the correct arm, followed by 5 free-choice trials. Correction within a trial was not permitted and the mice were returned to the start cylinder in a random orientation to prevent the adoption of motor response strategies (Figure 1E, C and Supplementary Figure 2).

To control for potential drug-related effects, a separate group of mice (n = 4 males) without DREADD expression in the medial septum was trained on the same task and received identical treatments. These control animals showed normal learning across training (Figure. 4A) and no difference in performance between DCZ and gelatin conditions (Figure. 4B). Working memory errors in control animals remained low and unchanged across treatments (Supplementary Figure 3A-B), confirming that the observed behavioural effects were specific to DREADD-mediated inhibition of MS function and not an ‘off-target’ effect of DCZ.

**Figure 4.**
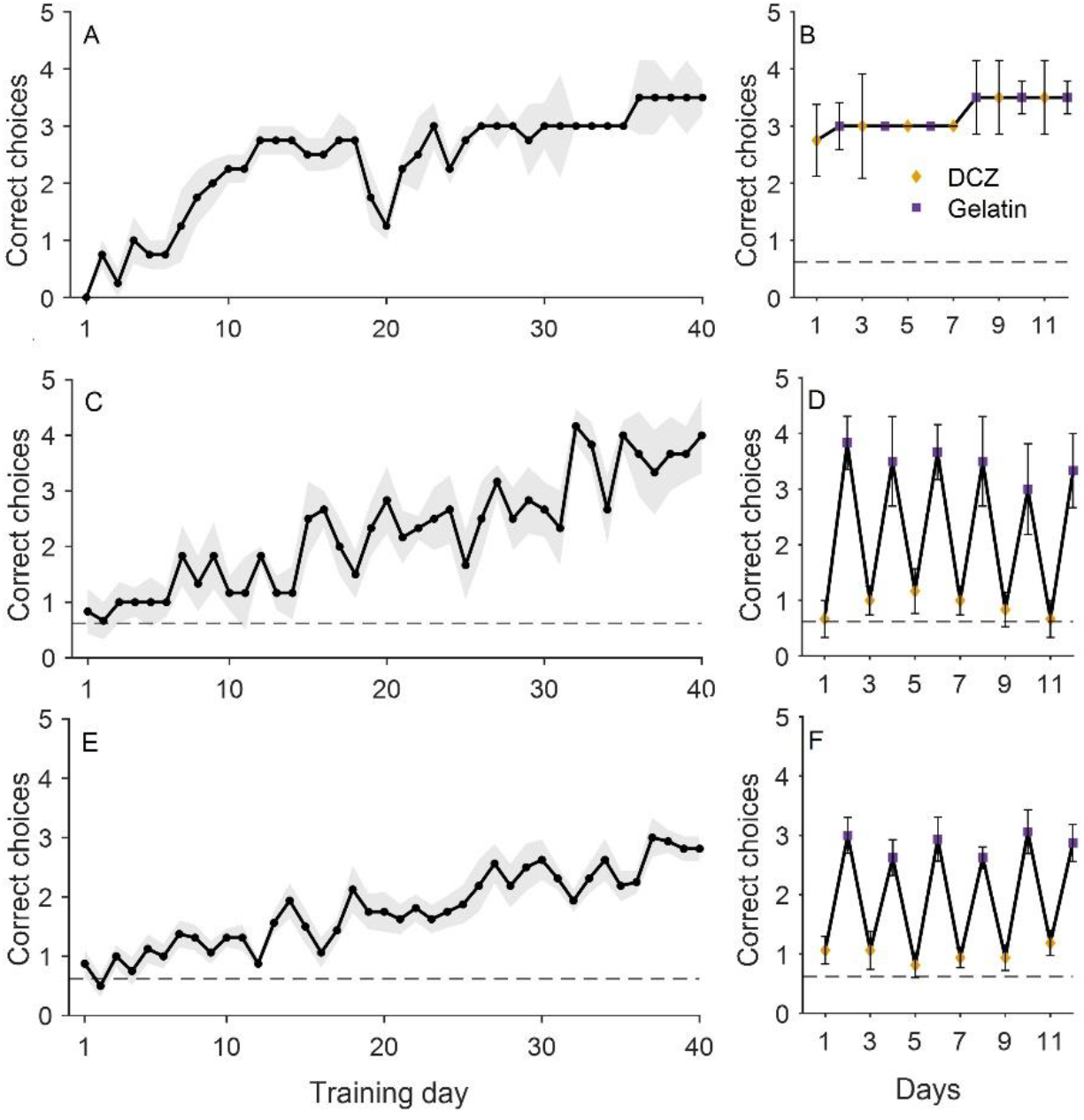
Performance across random reward-arm training and alternating DCZ/gelatin sessions. (A) Mean correct choices across 40 days of random reward arm training in the control male cohort (n = 4). The shaded region indicates SEM. (B) Mean correct choices during the 12 training sessions in the same control cohort, plotted as alternating DCZ and gelatin sessions. Orange diamonds indicate DCZ sessions and purple squares indicate gelatin sessions. Error bars indicate SEM. (C) Mean correct choices across 40 days of random reward arm training in the first male cohort (n = 6). (D) Mean correct choices during the 12 training sessions in the first male cohort, plotted as alternating DCZ and gelatin sessions (E) Mean correct choices across 40 days of random reward arm training in the second male cohort (n = 16). (F) Mean correct choices during the 12 training sessions in the second male cohort, plotted as alternating DCZ and gelatin sessions. Dashed horizontal lines indicate chance performance (0.625).

In Cohort 1, the male mice showed an increase in correct choices across days (Figure. 4C) and a decrease in working memory errors (Supplementary Figure 3C). Female mice showed relatively poorer learning across training days (Supplementary Figure 4A) barely exceeding chance performance and more variable working memory errors (Supplementary Figure 4C).

Following training, animals were tested over 12 days under alternating DCZ and gelatin conditions (6 DCZ days, 6 gelatin days). In male mice, DCZ severely reduced correct choices compared to gelatin days, with performance approaching chance levels (Figure. 4D). Working memory errors increased on DCZ days and were lower on gelatin days (Supplementary Figure 3D). In female mice, there was a similar decrement in correct choices under DCZ treatment although the control performance was poor (Supplementary Figure 4B). Working memory errors were also increased under DCZ (Supplementary Figure 4D). Together, these findings indicate that medial septum inhibition disrupts spatial memory performance under conditions that require daily updating of goal locations.

To quantify learning, performance was averaged across early (Days 1–10) and late (Days 31–40) phases. A two-way repeated-measures ANOVA revealed a significant effect of training phase (F(1,10) = 372.82, p < 0.001), a main effect of sex (F(1,10) = 7.23, p = 0.023), and a significant Sex × Phase interaction (F(1,10) = 15.26, p = 0.0029), indicating distinct learning dynamics between males and females. Because of the poor learning exhibited by female mice, the remaining studies were conducted exclusively with males. In the next study, a larger cohort of DREADD-expressing male mice (n = 16) was trained for 40 days on the same random reward-arm task until performance neared an asymptotic level (Figure. 4E). Animals were then tested under alternating DCZ and gelatin conditions. In this cohort, DCZ administration also consistently reduced performance, with correct choices decreasing toward chance levels (Figure. 4F), while performance on gelatin days remained stable and comparable to pre-treatment levels.

In addition, DCZ administration was associated with a modest increase in working-memory errors relative to gelatin days (Supplementary Figure 3E–F). Although overall error rates remained low throughout testing, a small increase was observed during the early DCZ sessions, reflected by more revisits to previously entered reward arms. These results further support a role for medial septal activity for within-session memory processes (Stackman & Walsh, 1994).

### With a fixed goal arm, performance gradually became resistant to MS manipulation

Following a 3-day break after completion of the random reward task, the male mice (n = 6) of Cohort 1 were trained on a fixed reward arm version of the task, in which the rewarded arm was randomly selected for each mouse but then remained constant across days. The animals were tested under alternating gelatin and DCZ conditions, starting with gelatin on the first day. As shown in Figure 5A, even on the first DCZ day (day 2), performance, although slightly reduced, was nevertheless substantially higher than on the random reward arm task under DCZ indicating that MS interneuron inhibition was less disruptive after having learned the correct arm on the previous day. Correct choices increased across testing days, indicating continued improvement in performance. Although performance on early DCZ days was lower relative to adjacent gelatin days, this difference diminished over time. By later sessions, performance remained high and comparable across DCZ and gelatin conditions, indicating reduced sensitivity to DREADD activation. In addition, working memory errors were low throughout training and showed a gradual decrease across days. While a slight increase in errors was observed during early DCZ sessions compared to gelatin days, this effect diminished with continued training. By later sessions, error rates were minimal and similar across both conditions (Supplementary Figure 5A). Together, these results indicate that, with repeated training on a fixed goal location, animals become progressively less dependent on normal medial septal function for both accurate choice behaviour and working memory performance.

**Figure 5.**
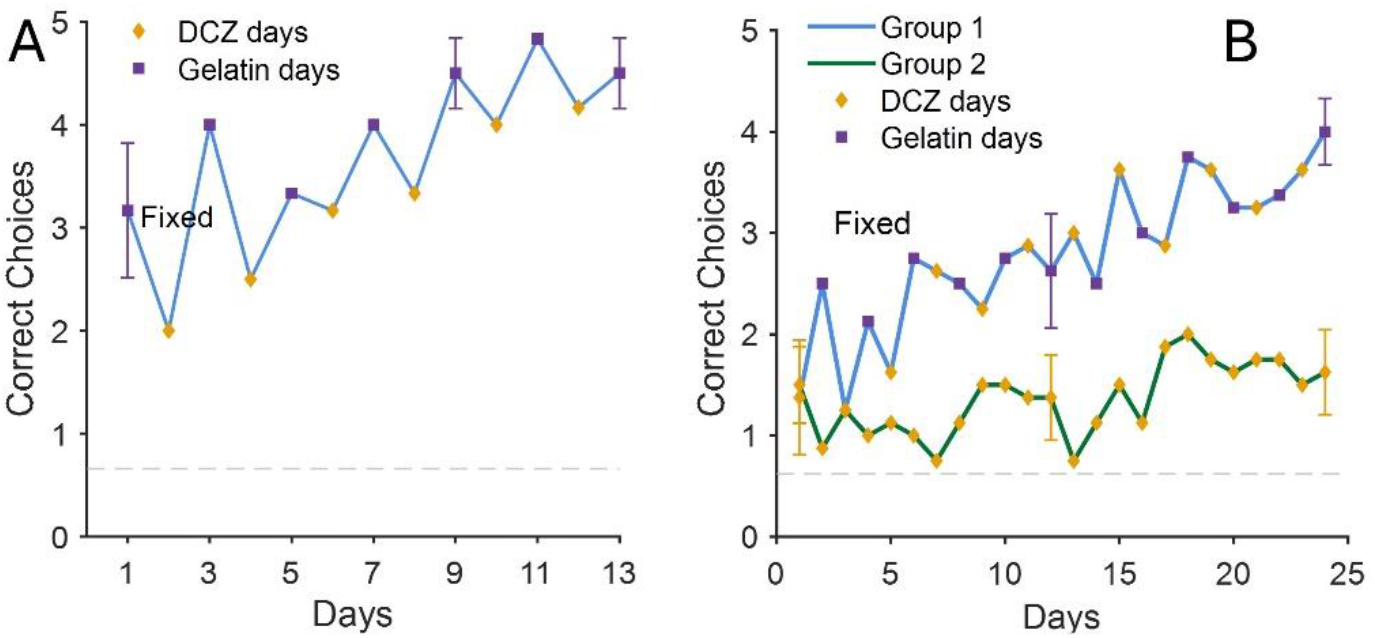
Performance during fixed reward-arm training and alternating DCZ/gelatin. (A) Mean correct choices across 13 days of fixed reward arm training in the first male cohort (n = 6), plotted as alternating DCZ and gelatin sessions. Orange diamonds indicate DCZ sessions and purple squares indicate gelatin sessions. Error bars indicate SEM. (B) Mean correct choices across 24 days of fixed reward arm in the second male cohort (n = 8 per group). Group 1 (blue) received alternating DCZ and gelatin sessions, and Group 2 (green) received daily DCZ. Orange diamonds indicate DCZ sessions and purple squares indicate gelatin sessions. Error bars indicate SEM. Dashed horizontal lines indicate chance performance (0.625).

To further examine this emerging resistance to medial septal inhibition, Cohort 2 mice (n = 16) were tested under the same fixed-reward condition in which the rewarded arm remained constant across sessions. Three days after the random reward arm tests, the animals were divided into two groups (n = 8/group): one group received daily DCZ, while the other received alternating DCZ and gelatin across 24 days of training (in this case, day 1 was a DCZ day). As in Cohort 1, in the alternating DCZ/gelatin group, there was a clear effect of DCZ on the first 2-3 days, after which DCZ no longer impaired performance. In contrast, in the daily DCZ group, performance remained substantially lower throughout training, showing only modest improvement over time (Figure 5B).

Working memory errors showed a complementary pattern. In the alternating-treatment group, errors gradually decreased across training, consistent with learning of the fixed reward location. In the daily DCZ group, error rates were higher and showed no improvement across sessions (Supplementary Figure 5B). Together, these results indicate that with extensive training, performance under fixed task conditions became insensitive to septal disruption when animals were given DCZ on alternate days, whereas there was little if any improvement over days in animals that were never exposed to the task without MS disruption.

### Fixed reward training resulted in a transition to MS-independent memory

To test whether prior training and/or familiarity with the environment alters sensitivity to medial septum manipulation, in the second cohort, the rewarded arm was rotated by 180° while remaining fixed across sessions for an additional 24 days. Treatment schedules were simultaneously switched between groups: mice previously receiving daily DCZ were reassigned to alternating DCZ and gelatin, whereas mice previously receiving alternating treatment were switched to daily DCZ. Under these conditions, both groups showed robust improvement across training (Figure 6A). Correct choices increased from near-chance levels at the start of reversal to high levels by the end of training, indicating successful acquisition of the new goal location. Notably, this improvement occurred in both groups, including animals receiving daily DCZ, suggesting that medial septum disruption no longer impaired learning under these conditions. Errors due to entries into the previously rewarded arm were initially near chance early in training and progressively declined across days as correct choices increased (Figure 6B).

**Figure 6.**
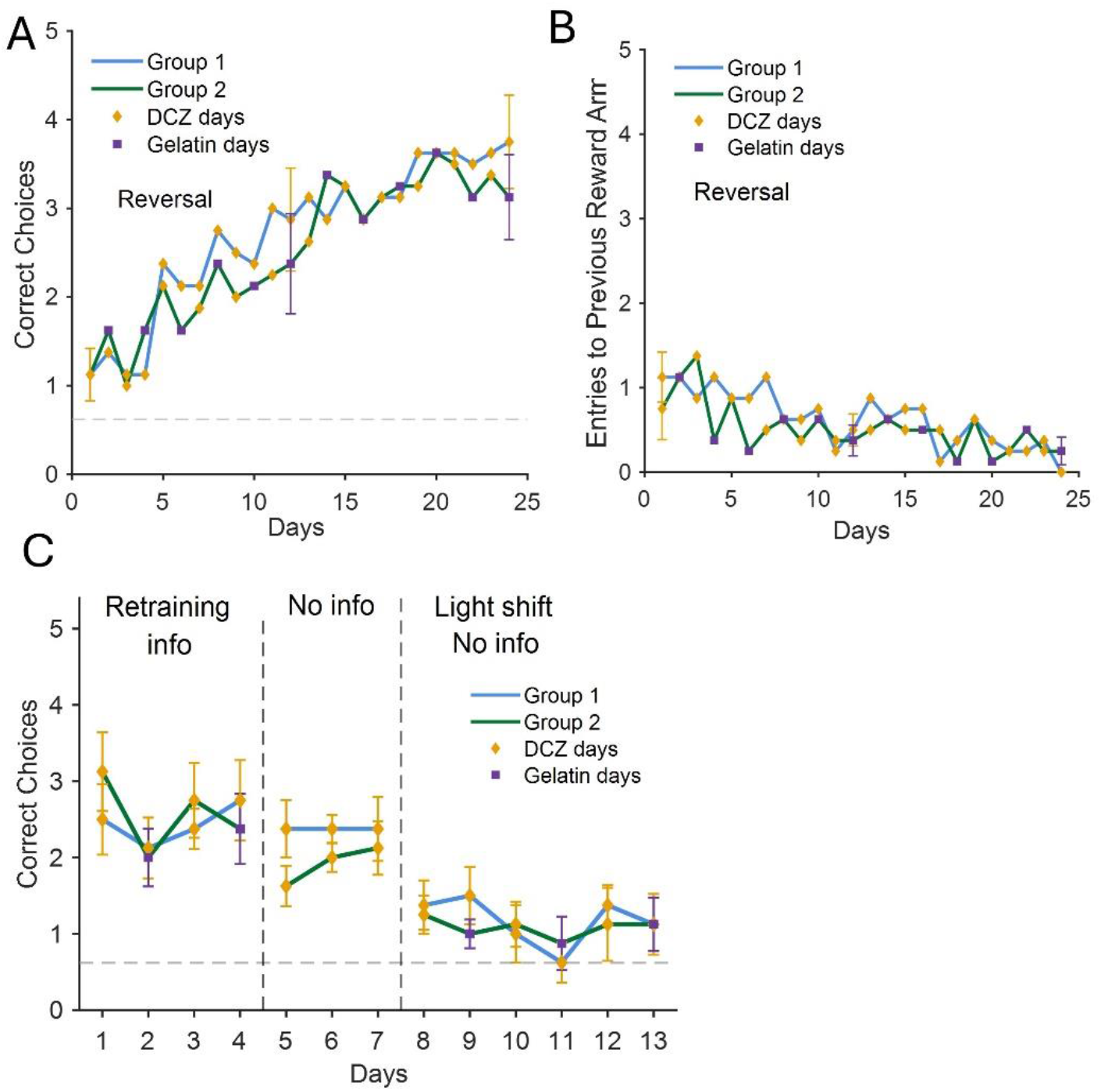
Performance during reversal learning and post-delay probe testing. (A) Mean correct choices across 24 days of reversal learning in the fixed reward-arm task (n = 8 per group). The rewarded arm was rotated 180° relative to the preceding fixed reward arm phase. Treatment schedules were switched between groups such that Group 1 received DCZ daily, whereas Group 2 received alternating DCZ and gelatin sessions. Orange diamonds indicate DCZ sessions, and purple squares indicate gelatin sessions. Error bars represent SEM for treatment-matched days. (B) Mean entries into the previously rewarded arm across 24 days of reversal learning (n = 8 per group). Group 1 and Group 2 are shown separately. (C) Mean correct choices during post-delay retraining and probe testing following a 3-week break (n = 8 per group). Sessions are divided into retraining, no-information trials, and light-shift phases, indicated by dashed vertical lines. Group 1 and Group 2 retained the same treatment schedules used during reversal learning. The dashed horizontal line indicates chance performance (0.625).

Working memory errors were elevated at the beginning of reversal training and declined across days, eventually converging across groups (Supplementary Figure 6 A). By the end of training, working memory performance was similar regardless of treatment conditions. Following Phase 2 (fixed reward arm), in which performance became resistant to medial septum interneuron inhibition, acquisition of a new fixed goal in the same environment in Phase 3 (reversal) was no longer impaired by DREADD activation. This generalization suggests that repeated fixed-goal training under MS-HPC axis disruption induced a reorganization of the neural systems supporting reward location memory in a familiar environment.

### Memory became resistant to septal inhibition but remained sensitive to global cue reconfiguration

To assess memory retention and cue dependence, animals were tested following a three-week break after completion of the reversal phase. Mice were first retrained for several days under the same fixed-reward configuration used in the reversal phase (Figure 6C, “Retraining +info”). Performance during retraining remained well above chance in both groups. Animals were then tested for several days without information trials (“No info”), during which all animals received DCZ and were required to rely on previously acquired spatial memory. Under these conditions, performance remained relatively stable, suggesting that animals were able to retrieve the learned reward location under the influence of MS GABAergic inhibition and in the absence of information trials.

To examine cue dependence, the position of the dominant distal cue in the room (light source) was rotated by 180° while the rewarded arm remained unchanged (“Light shift+no info”). Following this manipulation, correct choices were diminished across sessions in both groups (Figure 6C), accompanied by an increase in working memory errors (Supplementary Figure 6B). These changes were observed across both treatment groups. Instead, arm selections became more variable across sessions, indicating a disruption in task performance under the altered cue configuration.

To test the hypothesis that animals had been using the room light as a directional reference, arm selections following the light-shift manipulation were analyzed relative to the rotated cue configuration. Choices were expressed as angular deviations, where 0° corresponded to the correct (rewarded) arm and angles were defined relative to the shifted light position. This approach allowed assessment of whether animals followed the previously learned reward location or reoriented behaviour based on the displaced salient distal cue. The daily DCZ group exhibited a mean length of *r* = 0.05, whereas the alternating DCZ/gelatin group showed a mean vector length of *r* = 0.06. The short mean vectors in both groups indicated low directional consistency and high variability in arm selection following the light shift. Importantly, neither vector was oriented toward the direction predicted by the rotated light cue (i.e., 180° relative to the original configuration), indicating that animals did not simply follow the shifted light as a dominant directional reference. Instead, the absence of a clear directional bias suggests a disruption in the spatial information guiding behaviour, with a similar pattern observed in both treatment groups.

Together, these results indicate that the light-shift manipulation disrupted spatially guided behaviour in both groups, and that performance was not driven by a single salient cue. Rather, behaviour appeared to rely on a learned visual representation and was disrupted when the visual information was altered.

## Discussion

The electrophysiological results show that DREADD-mediated inhibition of medial septal interneurons reduced hippocampal theta power and significantly decreased the spatial information carried by hippocampal and cortical neurons while preserving overall neuronal firing rates. This dissociation between preserved firing rates and degraded spatial coding is consistent with prior work showing that septal disruption impairs the temporal structure of hippocampal activity without abolishing neuronal firing. Mizumori et al. (1989) demonstrated that reversible septal inactivation with lidocaine abolished theta and disrupted spatial memory performance while qualitatively preserving place-specific firing in CA1, and subsequent studies showed that theta disruption impaired grid cell periodicity and theta sequences without eliminating spatial selectivity (Brandon et al., 2011; Wang et al., 2015). Disruption of medial septal circuits with the GABAA antagonist Gabazine, impaired hippocampal theta and spatial navigation, even when substantial place-cell firing remained preserved (Bolding et al., 2020). Our findings extend this literature by showing that suppression of septal inhibitory interneurons reduced the information content of neural activity across both hippocampus and cortex, supporting the view that septal input is essential for maintaining the quality and structure of spatial representations. It is important to bear in mind that the effects of various septal manipulations may be somewhat dependent on exact experimental conditions such as task, experience, species, age etc.; however, the general conclusion that the fine temporal structure of hippocampal neuronal firing is important for normal spatial learning and memory appears sound. Previous work has further demonstrated that coordinated interactions between hippocampal, cortical, and thalamic networks are important for memory-related information processing and systems-level communication (Alizadeh et al., 2022; Azimi et al., 2021). The present findings raise the possibility that medial septal interneurons and septo-hippocampal theta mechanisms may play a critical role in organizing this large-scale coordination during spatial learning; however possible changes in synaptic plasticity mechanisms are not ruled out.

Consistent with this physiological result, we found that inhibition of MS interneurons impaired spatial learning in the radial arm maze task when animals were required to encode a randomly selected goal location each day. Under this condition, performance dropped to near chance and working memory errors increased, indicating a failure to acquire place-reward associations. This finding aligns with classic studies demonstrating that septal lesions or inactivation disrupt spatial learning and abolish hippocampal theta (Winson, 1978; Stewart and Fox, 1990), as well as with work linking septal input to synaptic plasticity and learning (Leung and Shen, 1995; Hölscher et al., 1997, Kubik et al., 2007). However, the contribution of different septal cell populations is not equivalent, as selective depletion of medial septal cholinergic neurons alone has been shown to place learning largely intact (Craig et al., 2008, 2009, 2011). Together with our physiological results, these data reinforce the conclusion that intact septo-hippocampal function is important when behaviour requires flexible encoding and rapid updating of place-reward associations.

In contrast, when the rewarded arm remained constant across days, the impact of septal disruption progressively diminished. Although performance was initially impaired, animals gradually became resistant to DREADD activation. This reduced behavioural sensitivity is unlikely to reflect a loss of DREADD efficacy resulting from repeated DCZ exposure. Prior work has shown that orally administered DCZ is an effective DREADD agonist for repeated non-invasive administration (Ferrari et al., 2022). Extending these findings, our pilot experiments revealed no reduction in the behavioural efficacy of DCZ across repeated administration over several months, with its behavioural effect persisting for at least 1 hour following administration (Supplementary Figure 7), indicating that the progressive loss o impairment observed here is more likely to reflect experience-dependent changes in the neural systems supporting task performance rather than pharmacological desensitization. Previous studies have shown that spatial firing can persist despite septal disruption (Mizumori et al., 1989; Brandon et al., 2014), but these studies examined acute effects and did not address how the behavioural consequences of septal disruption may change with learning. Our findings provide an extension by demonstrating that the behavioural reliance on septal input is dynamic and can be reduced through repeated experience.

A key finding of the present study is that the reduced dependence on MS generalized beyond the originally learned goal. After animals became resistant to septal inhibition in the fixed-goal condition, they were able to acquire a new goal location without impairment under MS inhibition. This observation goes beyond previous work by showing that, in some cases, training does not simply lead to overlearning of a specific memory but instead produces a more general change in the neural systems or strategies supporting behaviour. One possible interpretation is that repeated fixed-goal training promotes the formation of a schema or task representation that can support both retrieval and new learning within a stable environment. This idea is consistent with proposals that repeated experience allows animals to extract regularities and organize behaviour more efficiently (Sutherland et al., 2010; Sutherland and Lehmann, 2011). The novel contribution of our study is to link this systems-level reorganization to a reduced dependence on normal medial septum function during spatial behaviour.

These findings have important implications for theories of systems consolidation. Classical models propose that memories gradually become independent of the hippocampus over time (Squire and Zola-Morgan, 1991; Frankland and Bontempi, 2005), supported by evidence for increased cortical involvement during remote memory retrieval (Bontempi et al., 1999; Frankland et al., 2004). However, many rodent studies of spatial memory have challenged this view, reporting persistent hippocampal dependence even at remote time points (Sutherland et al., 2001; Martin et al., 2005; Clark et al., 2005). Our results suggest that this apparent discrepancy may arise because hippocampal dependence is influenced not only by time but also by the structure and content of training. Specifically, repeated experience with a stable goal relationship may allow animals to develop representations or strategies that reduce reliance on septal input and potentially on hippocampal processing more broadly.

A plausible interpretation of these findings is that, with extensive training in a familiar environment, animals may have shifted from a hippocampus-dependent learning strategy to a strategy based on visual pattern matching. Under this account, repeated experience would allow animals to associate a particular visual scene with the rewarded location. An alternative possibility is that repeated training to a fixed goal location allowed other parallel learning and memory systems to gradually acquire a place response. For instance, the dorsal striatum has been proposed to support place responses under repeated fixed-location training, providing a non-hippocampal solution to spatial tasks (McDonald and White, 1995). Consistent with this interpretation, performance was impaired when the visual configuration was altered by rotating the light (Supplementary Figure 8), suggesting that stable visual information had become an important determinant of behaviour. Such a strategy would not require normal hippocampal function. Previous studies have shown that hippocampal lesions do not impair the acquisition of simple visual discrimination tasks, whereas deficits emerge when successful performance requires relational representations, contextual processing, or flexible memory retrieval (Hales et al., 2015). Instead, visual discrimination and item-based learning rely primarily on visual cortical and perirhinal circuits, with perirhinal cortex becoming important (Mumby and Pinel, 1994; Kolb et al., 1994; Eacott et al., 2001). Similarly, hippocampal involvement in object recognition is generally limited to tasks with greater contextual or episodic memory demands rather than simple familiarity or visual pattern matching (Cohen and Stackman, 2015). Together, these findings raise the possibility that extensive fixed-goal training allowed animals to learn to solve the task by gradually associating specific visual configurations with reward. Alternatively, repeated fixed-goal training may have allowed other learning and memory systems to gradually acquire a solution to the task. Once these strategies were adopted, animals were able to learn a new configuration–reward association despite disruption of normal MS function. Consistent with evidence that hippocampal learning can occur more rapidly than learning supported by other systems (e.g., McClelland et al., 1995; McDonald and White, 1995), this learning was substantially slower than when normal septo-hippocampal function was available. An additional finding was a sex difference in acquisition. Females showed weaker improvement across training than males. The significant Sex × Phase interaction suggests sex-dependent differences in learning across training. However, because subsequent experiments included only males, the effects of medial septal GABAergic inhibition should not be assumed to generalize to females.

In summary, our results show that medial septal interneurons are critical for the initial acquisition of place-reward associations under conditions requiring flexible updating, but that this dependence can be reduced through repeated experience in a stable environment. By demonstrating that septal inhibition disrupts spatial coding without abolishing neuronal firing, and that its behavioural consequences change with learning, this study provides new evidence that the contribution of the medial septum to behaviour is dynamic rather than fixed. The observed deficits may reflect disruption of septo-hippocampal network dynamics, reduced support for hippocampal plasticity, or a combination of both mechanisms. Alternatively, repeated training may promote a transition from a hippocampus-dependent learning strategy to one based on gradually learned visual pattern matching or visual configuration recognition, reducing reliance on normal medial septum function. Distinguishing between these possibilities will require future experiments that directly dissociate spatial navigation from visual cue-based strategies.

## Materials and Methods

### Subjects

#### Electrophysiological recordings

Two adult male wild-type mice (B6129SF1/J) were used. Mice were approximately 3 months of age at the time of surgery and were singly housed with food and water available *ad libitum*. Animals underwent one week of handling prior to surgery. Food restriction was initiated one week after surgery, before recording sessions.

#### Behavioural experiments

A total of 32 Thy1-GCaMP mice were used, comprising three experimental groups (cohort 1: n = 6 males and n = 6 females, cohort 2: n = 16 males) and a control group (n = 4) that did not undergo surgical procedures or viral injections; all animals were 42 days old at the start of the experiment. Mice were singly housed with food and water available *ad libitum* and underwent one week of handling prior to surgery. Food restriction (not more than 85% reduction in *ad libitum* weight) was initiated at the onset of radial arm maze training, at which point animals were approximately 2.5 months of age.

All animals were maintained at 24°C under a 12 h light/dark cycle (lights on at 7:30 a.m.), and all testing was conducted during the light phase. All procedures were approved by the appropriate institutional animal care committees at both the University of Lethbridge and the University of California, Irvine, and were carried out in accordance with national guidelines for the care and use of laboratory animals. Female mice were excluded because prior data indicated that they did not learn this task as reliably as males (Supplementary Figure 4).

### Surgical Procedures

Mice underwent stereotaxic surgery for viral delivery of inhibitory DREADDs to the medial septum (MS) using the construct pAAV-hDlx-GiDREADD-dTomato-Fishell-5 (Addgene, Watertown, MA). All procedures were performed under aseptic conditions.

For behavioural experiments, animals received buprenorphine (0.1 mg/kg, subcutaneous) 30 minutes prior to surgery for preoperative analgesia. Anesthesia was induced with isoflurane (3%) and maintained at 1.0–1.5% in oxygen (1L/min). Mice were placed in a stereotaxic frame, and body temperature was maintained at 37°C using a feedback-controlled heating pad. Animals also received dexamethasone (2 mg/kg, intramuscular) to minimize inflammation and saline (1 mL, subcutaneous) for hydration.

Lidocaine (7 mg/kg, subcutaneous) was applied locally at the incision site. The scalp was shaved, disinfected with chlorhexidine-alcohol, and incised along the midline to expose the skull. A burr hole was drilled above the medial septum, and the viral vector was delivered using a Hamilton syringe connected to a microinfusion pump through a 32-gauge cannula (Harvard Apparatus). A total volume of 0.4 microliters was injected at the following stereotaxic coordinates relative to bregma: anterior-posterior (AP), +1.4 mm; medial-lateral (ML), 0.0 mm; dorsal-ventral (DV), 4.3 mm. Injections were performed at a rate of 0.15 microliter per min. After infusion, the cannula was left in place for 5 minutes to allow diffusion and minimize backflow before being slowly withdrawn. Minor sinus bleeding after cannula withdrawal was controlled with gelfoam.

For electrophysiological experiments, animals underwent a similar stereotaxic procedure under isoflurane anesthesia (induction at 3%, maintenance at 1.5%), with additional administration of diazepam (5 mg/kg, intraperitoneal) and meloxicam (1–2 mg/kg, subcutaneous) for sedation and analgesia. Atropine (0.1 mg/kg, subcutaneous) was administered to reduce respiratory secretions. Viral injections were performed in the medial septum at AP +1.2 mm, ML 0.0 mm, and two depths (DV 4.5 mm, 0.25 microliter; DV 4.4 mm, 0.15 microliter) using a 32-gauge cannula attached to a microinfusion pump (0.15 microliter per min). The cannula was left in place for 3 minutes after each injection and, for the second injection, was retracted by 0.5 mm and held for an additional 10 minutes before removal to ensure diffusion. For head fixation, two thin metal bars (6 mm × 4 mm × 0.5 mm) were attached bilaterally above the ears using dental cement (Metabond). The exposed skull was sealed with Kwik-Sil followed by additional dental cement. Following surgery, animals used for electrophysiological recordings received extended-release buprenorphine (Ethiqa-XR, 3.25 mg/kg). All animals were placed in a temperature-controlled recovery chamber until fully recovered. Post-operative care was administered for 3 days and included meloxicam (1 mg/kg, subcutaneous) and saline (1 mL, subcutaneous). Animals were allowed to recover for approximately 4 weeks to ensure adequate viral expression prior to subsequent experiments.

### Electrophysiology section

#### Craniotomy and catheterization

The day prior to electrophysiological recording, mice were anesthetized with isoflurane and the dental cement (Metabond) and Kwik-Cast covering the skull were removed. A craniotomy (3 mm diameter) was made over the target recording area, and the dura mater was carefully removed using a sterile 27G needle fashioned into a hook, and fine forceps. Any bleeding was controlled using Gelfoam. Following dural removal, the craniotomy was filled with sterile silicone gel (Dow Corning) to prevent dehydration and protect the cortical surface. The site was then sealed with Kwik-Cast and reinforced with Metabond. During the same procedure, a subcutaneous catheter (flexible tubing) was implanted in the back of the animal and secured to the headcap to allow administration of deschloroclozapine (DCZ) during recording sessions while the animal was head-fixed. At the end of the procedure, animals received meloxicam (5 mg/kg, subcutaneous) for postoperative analgesia.

#### Apparatus

The experimental setup consisted of a head-fixed virtual reality (VR) system combined with electrophysiological recording. Mice were positioned on a spherical treadmill (running ball) while head-fixed using a custom headfork. The ball was coupled to a rotary encoder, which continuously updated the animal’s position in a circular VR track (150 cm per lap). Visual stimuli were presented using three 12.4x 21 cm tablets (Galaxy Tab4, T330NU, Samsung) positioned at left, right and center positions relative to the animal, providing a 300° visual field coverage. The VR environment consisted of a desert-like context with spatial cues distributed along the track. A lick spout equipped with a sensor was positioned in front of the animal to detect licking behaviour. Rewards (15% condensed milk solution) were delivered via a reward-dispensing metal spout operated by a solenoid valve controlled by a custom-made MATLAB Graphical User Interface (GUI). The MATLAB software-controlled reward delivery, recorded the animal’s position in the virtual reality (VR) environment (at a rate of 30 Hz), and the timing of licking events, brakes were applied to the treadmill during the intertrial intervals between running laps. Additionally, digital synchronization pulses were transmitted to the acquisition system whenever the animal moved, licked, or the brakes were applied.

#### Behavioural training and task

Following recovery from surgery, during which inhibitory DREADDs were infused into the medial septum and head bars were implanted, animals were handled daily and placed on a food-restriction schedule until they reached approximately 85% of their ad libitum body weight. Mice were then gradually habituated to head restraint and trained 5 days per week for approximately 11–18 weeks before recording. One complete lap was approximately 150 cm in length, and animals were trained to traverse the full track to obtain a food reward delivered at the end of each lap. Session duration was progressively increased until animals could perform for up to approximately 1 h per day. A lick spout positioned in front of the mouth delivered reward at the end of the track, and animals learned to reliably lick the spout during reward delivery.

#### Silicon probe recording

The day after the craniotomy, mice were mounted in the recording apparatus, and the Metabond and Kwik-Cast were removed to expose the craniotomy. Silicon microprobes coated with fluorescent dye were lowered to stereotaxically defined coordinates using a micromanipulator (MP-285, Sutter Instrument), with insertion monitored under a surgical microscope (Zeiss OPMI Pico). Electrophysiological signals were acquired using Intan RHX headstages, which amplified, sampled (30 kHz), digitized, and filtered local field potentials (LFPs) from 256 channels. After the probe reached the target depth, a drop of mineral oil was applied to the exposed cortical surface, followed by a 45-minute settling period before recordings began. At the end of the experiment, the probe was slowly withdrawn, and the mouse was euthanized according to standard procedures.

#### Recording protocol

Recordings were performed only after mice reached stable performance in the virtual reality (VR) task and consistently completed sufficient laps to generate robust hippocampal theta during locomotion. Each recording session began with a ∼30 min baseline period in the familiar VR environment, during which animals ran repeated laps to obtain reward at the end of the track. Baseline recordings were followed by a subcutaneous injection of DCZ (0.5 mg/kg) and approximately 60 min of continued recording. Neural activity was recorded continuously before and after DCZ administration, allowing direct comparison of hippocampal network dynamics under baseline conditions and during acute medial septum inhibition. DREADD activation was performed only during recording sessions to assess its immediate effects on hippocampal oscillatory activity and unit firing under stable behavioural conditions.

##### Analysis

Local field potential (LFP) and spiking analyses were performed in MATLAB (MathWorks). Time–frequency analysis of the LFP was computed using MATLAB’s spectrogram function with a 2 s Hanning window and 50% overlap. Oscillatory power in the theta band (5–10 Hz) was quantified relative to delta-band power (1–4 Hz) to assess changes in hippocampal rhythmic activity following medial septal inhibition. To evaluate the effects of DREADD activation on neuronal spiking, firing rates were quantified for both multi-unit activity (MUA) and isolated single units. Pre- and post-DCZ firing rates were calculated using 5 min analysis windows. The pre-DCZ epoch was defined as the 5 min immediately preceding DCZ administration, whereas the post-DCZ epoch began 5 min after administration to allow sufficient time for drug onset. These same pre- and post-DCZ windows were used to quantify spatial information and temporal firing structure. Spatial information was calculated using the Skaggs information metric (Skaggs et al., 1993), and temporal firing structure was assessed using autocorrelograms of unit spike times. Statistical comparisons between pre- and post-DCZ epochs were performed using paired t-tests. For spatial coding analyses, linearized rate maps were computed by dividing the virtual track into 3 cm spatial bins and calculating firing rate within each bin. Rate maps were then smoothed with a Gaussian kernel (1-bin standard deviation) prior to further analysis.

### Behavioural section

#### Drug administration

To habituate animals for oral delivery, mice received gelatin alone in their home cages for three days prior to testing. Following habituation, DCZ was administered orally via gelatin pellets in the home cage 10 minutes before behavioural testing.

Gelatin was prepared by dissolving 7 g unflavored gelatin (Knox) and 10 g Splenda (No Calorie Sweetener, with sucralose) in 49 mL distilled water, followed by the addition of 1 ml natural strawberry flavor. The mixture was heated to 50°C and stirred until fully dissolved. Aliquots (0.15 ml) were dispensed onto Falcon tube caps to form individual pellets (approximately 1 cm diameter and 1mm thick) and stored at 4°C. For DCZ administration, animals were weighed across three consecutive days, and the average body weight was used to determine the dosage. DCZ (5 mg/kg) was prepared from a 1 mg/mL stock solution and incorporated into gelatin immediately prior to pellet formation. Control animals received identical gelatin pellets without DCZ.

#### Radial arm maze procedure

Mice were food-restricted to approximately 85% of their body weight prior to behavioural testing. Behavioural testing was performed on an 8-arm radial maze (Figure 1). Each daily session consisted of two information trials followed by five free-choice test trials. During each information trial, the animal was placed at the center of the maze inside an opaque cylinder with an opening aligned with the rewarded arm, thereby guiding it toward that arm. On these trials, the mice were placed facing 180 deg opposite the gate. Once the animal entered the rewarded arm, the cylinder was rotated to confine the animal within the arm. After consuming the reward, the mouse was allowed to explore the rewarded arm for 30 seconds. The animal was then removed and returned to the center of the maze inside an opaque cylinder without an opening, with a piece of duct tape inside the cylinder consistently oriented toward the light direction. After 30 seconds, the cylinder was lifted, allowing the animal to choose an arm. During the five free-choice trials, correct choices were rewarded, after which the animal was allowed to consume the reward, picked up, and returned to the center of the maze inside the closed cylinder to begin the next trial. If the animal made an incorrect choice, it was immediately removed and returned to the center to initiate the next trial without receiving a reward.

#### Phase 0 (Initial task acquisition)

Mice were trained for 40 days on the 8-arm radial maze with a randomly changing rewarded arm each day, requiring rapid encoding of a new goal location. Each session consisted of two information trials followed by five free-choice trials.

#### Phase 1 (Random-reward testing under alternating drug conditions)

Mice were tested for 12 days on the random-reward task with alternating oral DCZ (5 mg/kg) and gelatin. This within-subject design assessed the effect of medial septum inhibition during rapid spatial encoding.

All experimental and control mice completed Phase 0 and Phase 1.

#### Phase 2 (Fixed-reward training)

Mice were trained with a fixed rewarded arm for 17 days (cohort 1) or 24 days (cohort 2). During the 17-day phase, animals received alternating DCZ and gelatin treatments. During the 24-day phase, mice were rank-ordered based on their performance during the 12-day random-reward phase and then alternatingly randomly assigned in descending rank order, to one of two groups, so that the two groups were approximately equated on initial performance. The two groups consisted of an alternating DCZ/gelatin group and a daily DCZ group.

#### Phase 3 (Fixed-reward training after reward rotation and treatment switch)

Mice were trained for 24 days with the rewarded arm shifted 180° from its Phase 2 location. Treatment conditions were switched between groups to assess how treatment influenced learning of the new reward location.

#### Phase 4 (Retest after a three-week break and cue manipulation)

Following a three-week break, animals were retrained for four days using the same fixed reward-arm configuration established in Phase 3 and under the same group and treatment conditions including the information trials. After retraining, mice were tested for three days on the fixed-reward task without information trials to assess whether they could retrieve the learned reward location from long-term memory. Subsequently, the position of the dominant light cue in the room was rotated by 180°, and the no-information-trial condition was repeated. This manipulation was used to determine the extent to which performance depended on the spatial relationship between the reward location and this dominant distal cue and/or whether a substantial alteration in the visual appearance of the arms from the start cylinder was important.

Two cohorts (n = 6 and n = 16 males) completed Phase 2, whereas only the larger cohort (n = 16) proceeded to Phases 3–4. During these phases, after completion of the information trials, each mouse was removed from the maze and placed in a recovery cage in the same room while the maze was rotated and cleaned. The animal was then returned to the maze to complete the five free-choice trials. This procedure was implemented to minimize the contribution of odour cues.

#### Analysis and statistical tests

All analyses, data visualization, and statistical tests were performed in MATLAB (R2022b, MathWorks). Repeated-measures ANOVA was used to evaluate the effects of experimental conditions and training across sessions.

### Histology

Upon completion of the experiments, animals were deeply anaesthetized with isoflurane and transcardially perfused with phosphate-buffered saline (PBS), followed by fixative (4% paraformaldehyde (PFA) in PBS for behavioural experiments or 10% formalin for electrophysiological experiments). Brains were extracted and post-fixed overnight at 4°C, then cryoprotected in 30% sucrose in PBS containing 0.02% sodium azide. Coronal sections (40 micrometer) were obtained using either a freezing microtome (behavioural cohort) or a cryostat (electrophysiological cohort). For the behavioural cohort, sections were mounted onto charged Superfrost Plus slides (VWR), coverslipped with Vectashield H-1000 mounting medium (Vector Laboratories), sealed with nail polish, and imaged using a NanoZoomer slide scanner (Hamamatsu) with a 40× objective. For the electrophysiological cohort, sections were stained with DAPI and imaged using a fluorescence microscope (Keyence) to verify probe placement and DREADD expression.

## Acknowledgements

We thank Dr. Robert Sutherland for helpful discussions of the Cohort 1 data and valuable suggestions regarding the design of Phase 2 of Cohort 2. Dr. Boyer Winters for reviewing the manuscript and providing valuable feedback, and Dr. Winters and members of his laboratory for training in viral injection and surgical procedures. We also thank Dr. Behroo Mirza Agha, Di Shao, the animal care staff, and members of the Animal Welfare Committees at the University of Lethbridge and the University of California, Irvine, for their support in facilitating the animal experiments. We thank Tarlan Goharian and Irish Supleo for their assistance with animal training. This work was supported by the Natural Sciences and Engineering Research Council of Canada (NSERC; RGPIN-2023-04402, awarded to B.L.M.; RGPIN-2020-06929, awarded to R.J.M.), the Canadian Institutes of Health Research (CIHR; PJT-173523), and the National Institutes of Health (NIH), USA (PHS grant RF1NS132041), awarded to B.L.M., as well as an Alberta Innovates Graduate Student Scholarship and the NSERC CREATE TrainBioData Training Program, awarded to Z.A.

## Supplementary materials

**Supplementary Figure 1.**
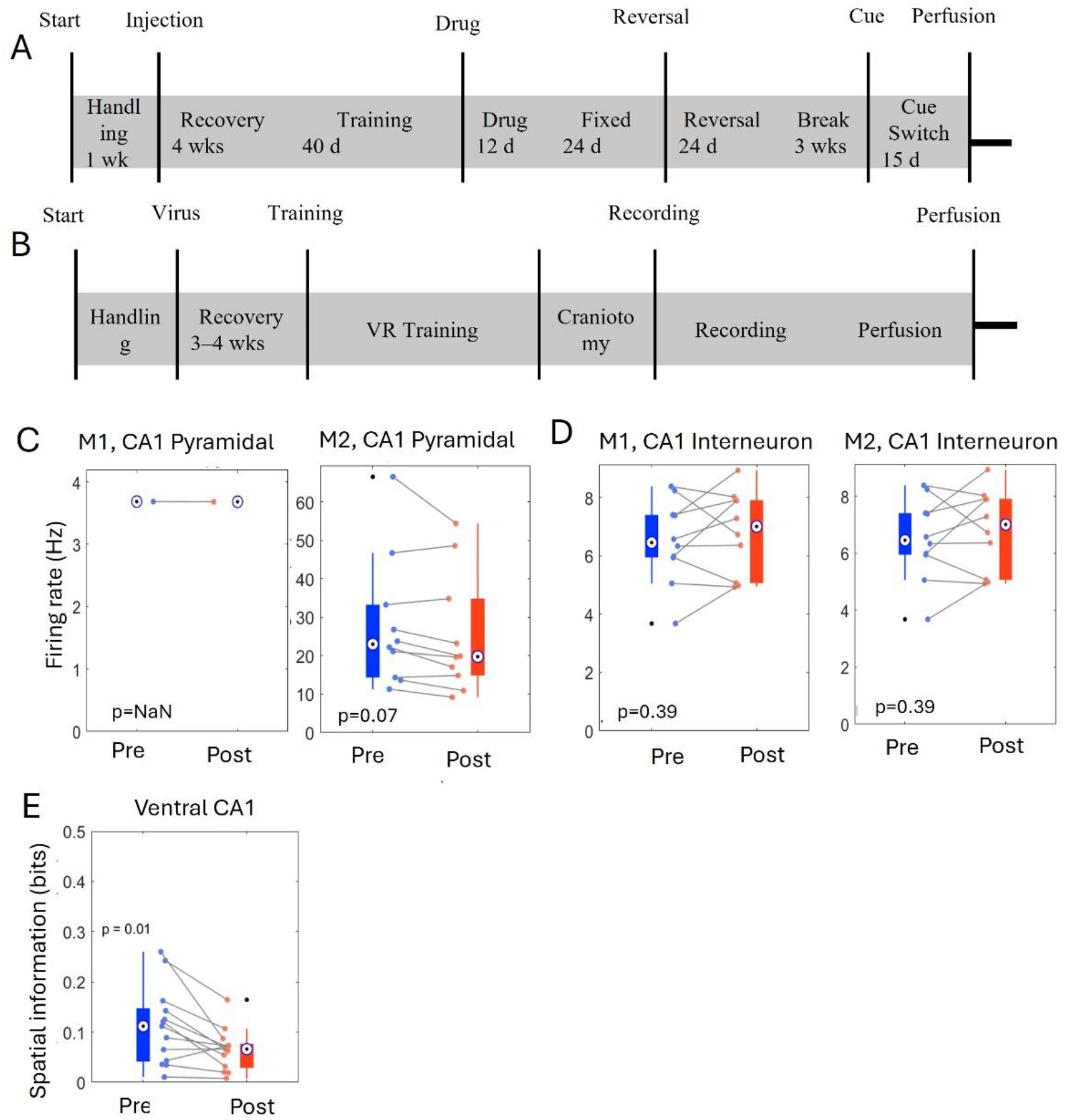
Experimental timelines and effects of medial septal inhibition on ventral CA1 firing and spatial information. (A) Behavioral timeline. Mice underwent 1 week of handling, 4 weeks of post-surgical recovery, 40 days of radial maze training, 12 days of alternating drug treatment, 24 days of fixed reward-arm training, 24 days of reversal learning, a 3-week break, and 15 days of post-delay cue-switch testing. (B) Electrophysiological timeline. Mice underwent handling, 3–4 weeks of post-surgical recovery, virtual reality (VR) training, craniotomy, recording sessions, and perfusion. (C) Firing rates of ventral CA1 pyramidal cells recorded before (Pre, blue) and after (Post, red) DCZ administration in two representative mice (Mouse1 and Mouse2). Each point represents one unit, with paired lines connecting pre- and post-DCZ values for the same cell. Boxplots show the distribution, median, and mean firing rate across units. (D) Firing rates of ventral CA1 interneurons recorded before (blue) and after DCZ administration (red) in the same two mice. (E) Spatial information of ventral CA1 pyramidal cells before (blue) and after DCZ administration (red).

**Supplementary Figure 2.**
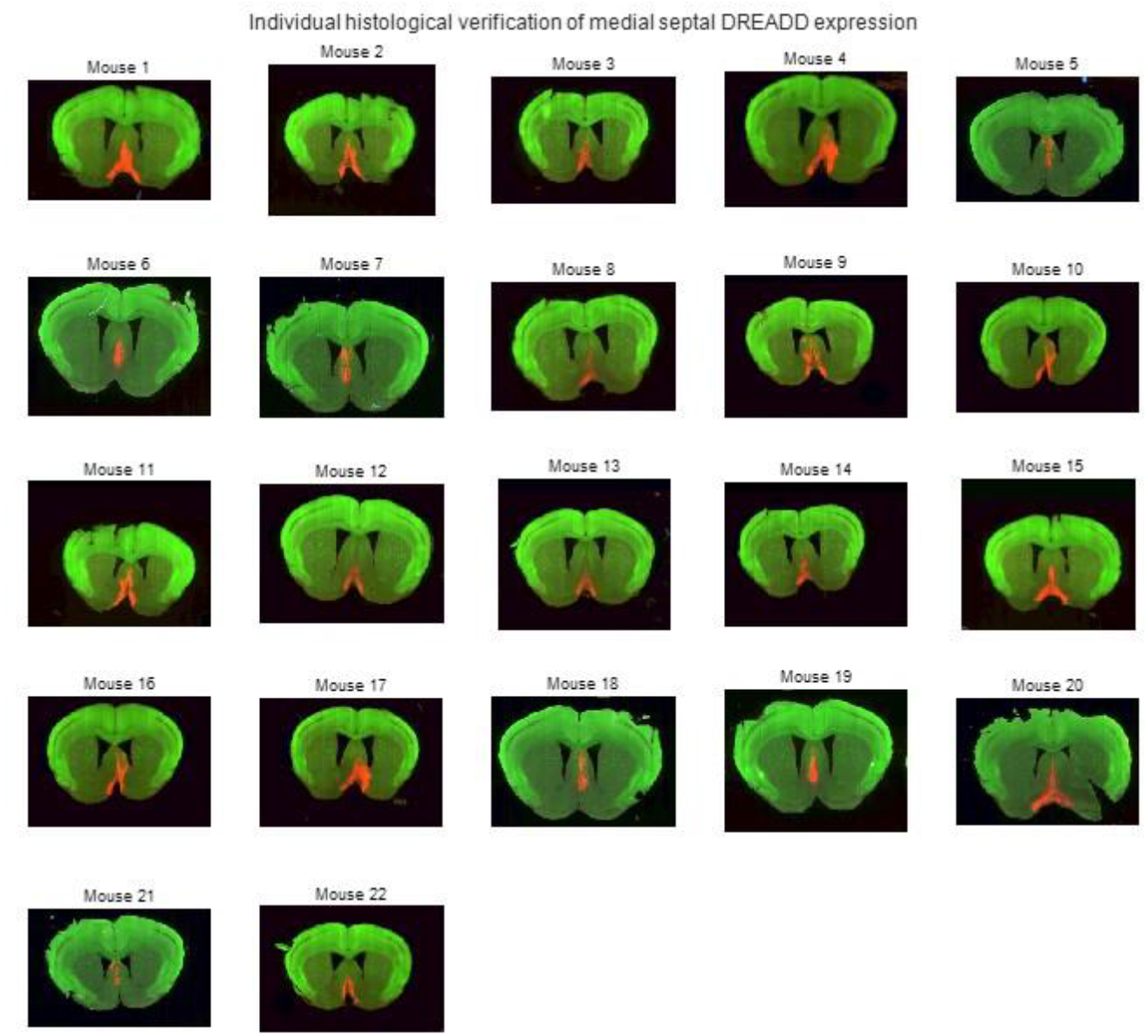
Histological verification of medial septal DREADD expression in individual animals. Representative coronal brain sections from individual mice included in the behavioral experiments, showing viral expression in the medial septal region. Green indicates tissue autofluorescence, and red indicates DREADD-associated viral expression. One representative section is shown for each animal (n=22).

**Supplementary Figure 3.**
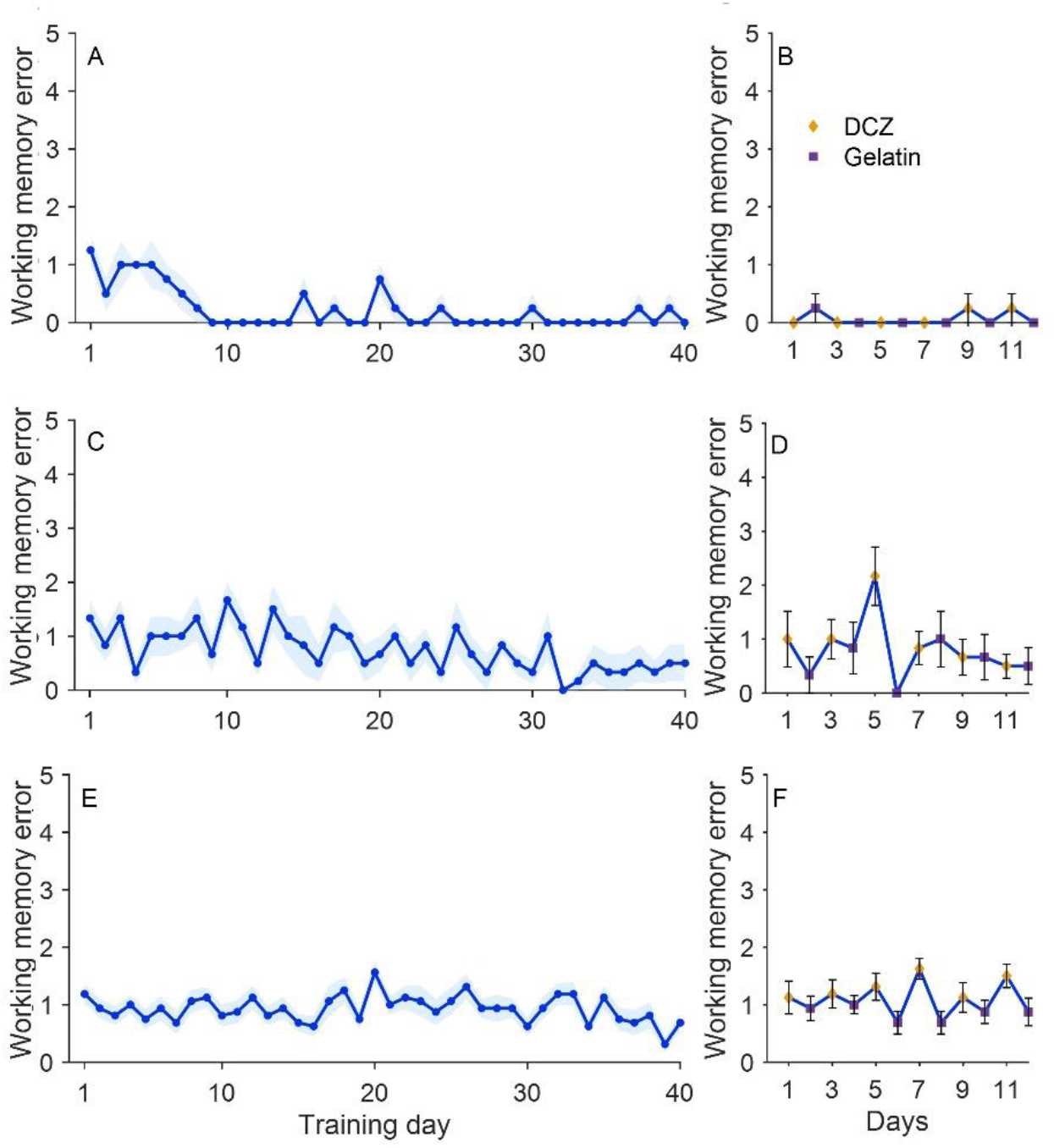
Working memory errors across random reward-arm training. (A) Mean working memory errors across 40 days of random reward-arm training in the control male cohort (n = 4). (B) Mean working memory errors across 12 alternating DCZ and gelatin sessions in the control male cohort (n = 4). (C) Mean working memory errors across 40 days of random reward-arm training in the first male cohort (n = 6). (D) Mean working memory errors across 12 alternating DCZ and gelatin sessions in the first male cohort (n = 6). (E) Mean working memory errors across 40 days of random reward-arm training in the second male cohort (n = 16). (F) Mean working memory errors across 12 alternating DCZ and gelatin sessions in the second male cohort (n = 16). In all figures, orange diamonds indicate DCZ sessions and purple squares indicate gelatin sessions. Error bars represent SEM.

**Supplementary Figure 4.**
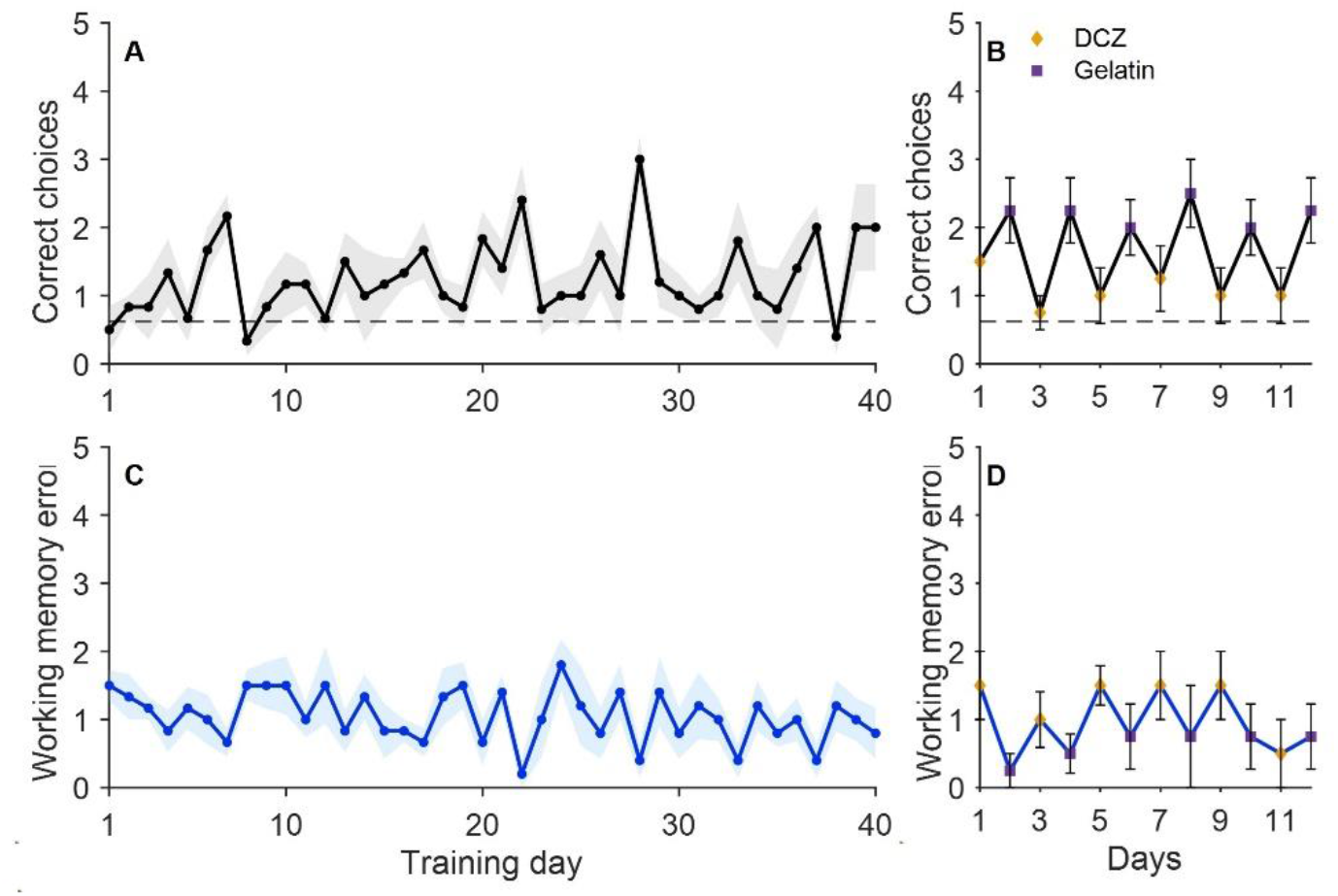
Random reward-arm performance. (A) Mean correct choices across 40 days of random reward-arm training in female mice (n = 6). (B) Mean correct choices across 12 alternating DCZ and gelatin sessions in female mice (n = 6). Orange diamonds indicate DCZ sessions and purple squares indicate gelatin sessions. Error bars represent SEM. The dashed horizontal line indicates chance performance (0.625). (C) Mean working memory errors across 40 days of random reward-arm training in female mice (n = 6). (D) Mean working memory errors across 12 alternating DCZ and gelatin sessions in female mice (n = 6).

**Supplementary Figure 5.**
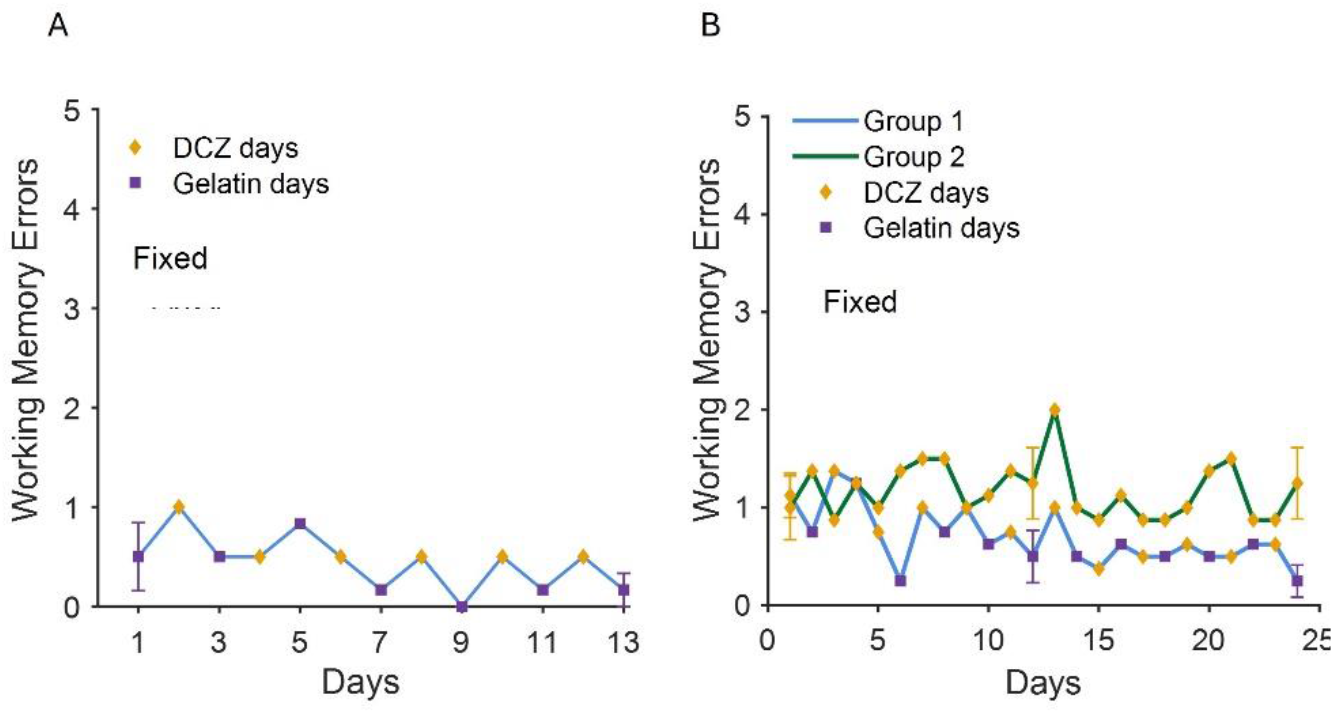
Working memory errors during fixed reward-arm training. (A) Mean working memory errors across 13 days of fixed reward-arm training in the first male cohort (n = 6). Orange diamonds indicate DCZ sessions and purple squares indicate gelatin sessions. Error bars represent SEM for treatment-matched days. (B) Mean working memory errors across 24 days of fixed reward-arm training in the second male cohort (n = 8 per group), shown separately for Group 1 and Group 2.

**Supplementary Figure 6.**
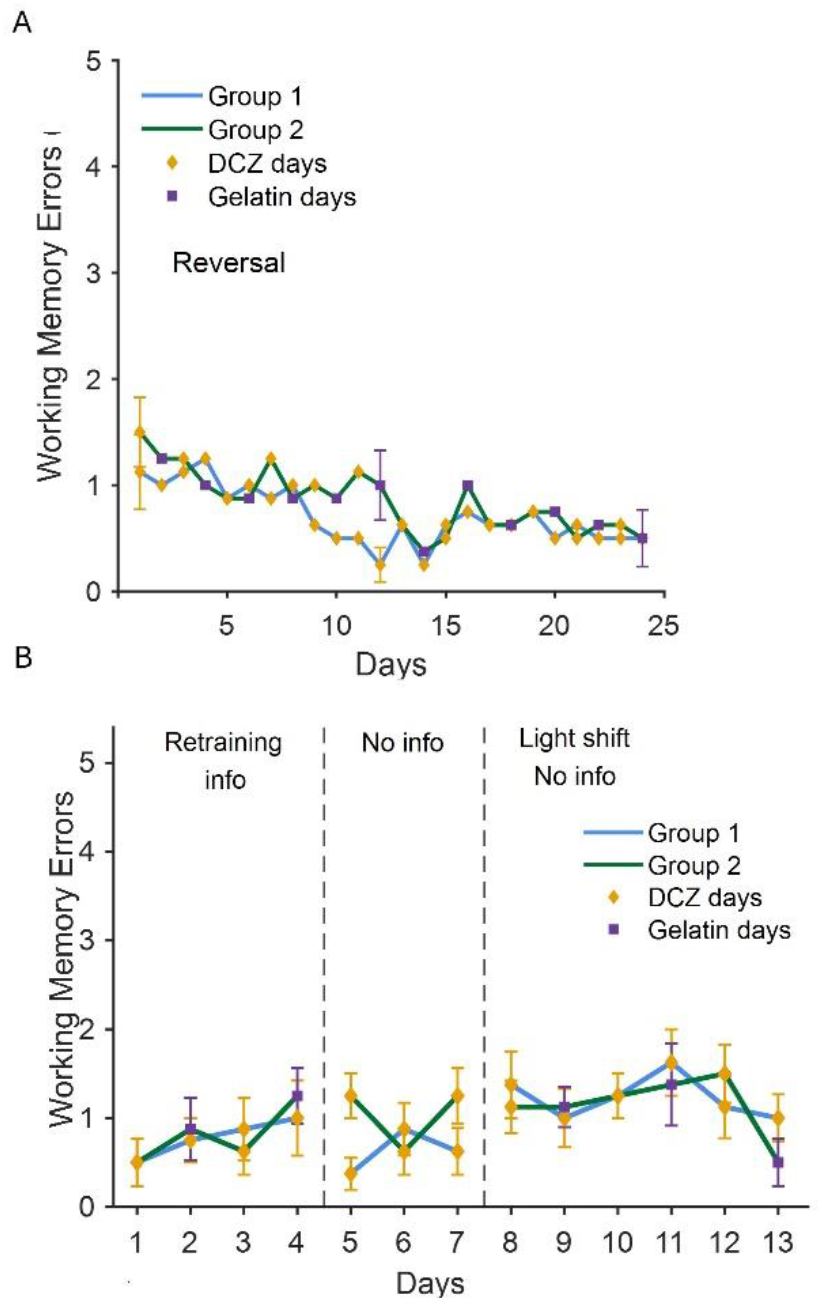
Working memory errors during reversal learning and post-delay probe testing. (A) Mean working memory errors across 24 days of reversal learning in the fixed reward-arm task (n = 8 per group). Group 1 and Group 2 are shown separately. Orange diamonds indicate DCZ sessions and purple squares indicate gelatin sessions. Error bars represent SEM for treatment-matched days. (B) Mean working memory errors during post-delay retraining and probe testing following a 3-week break (n = 8 per group). Sessions are divided into retraining, no-information, and light-shift phases, indicated by dashed vertical lines. Group 1 and Group 2 are shown separately.

**Supplementary Figure 7.**
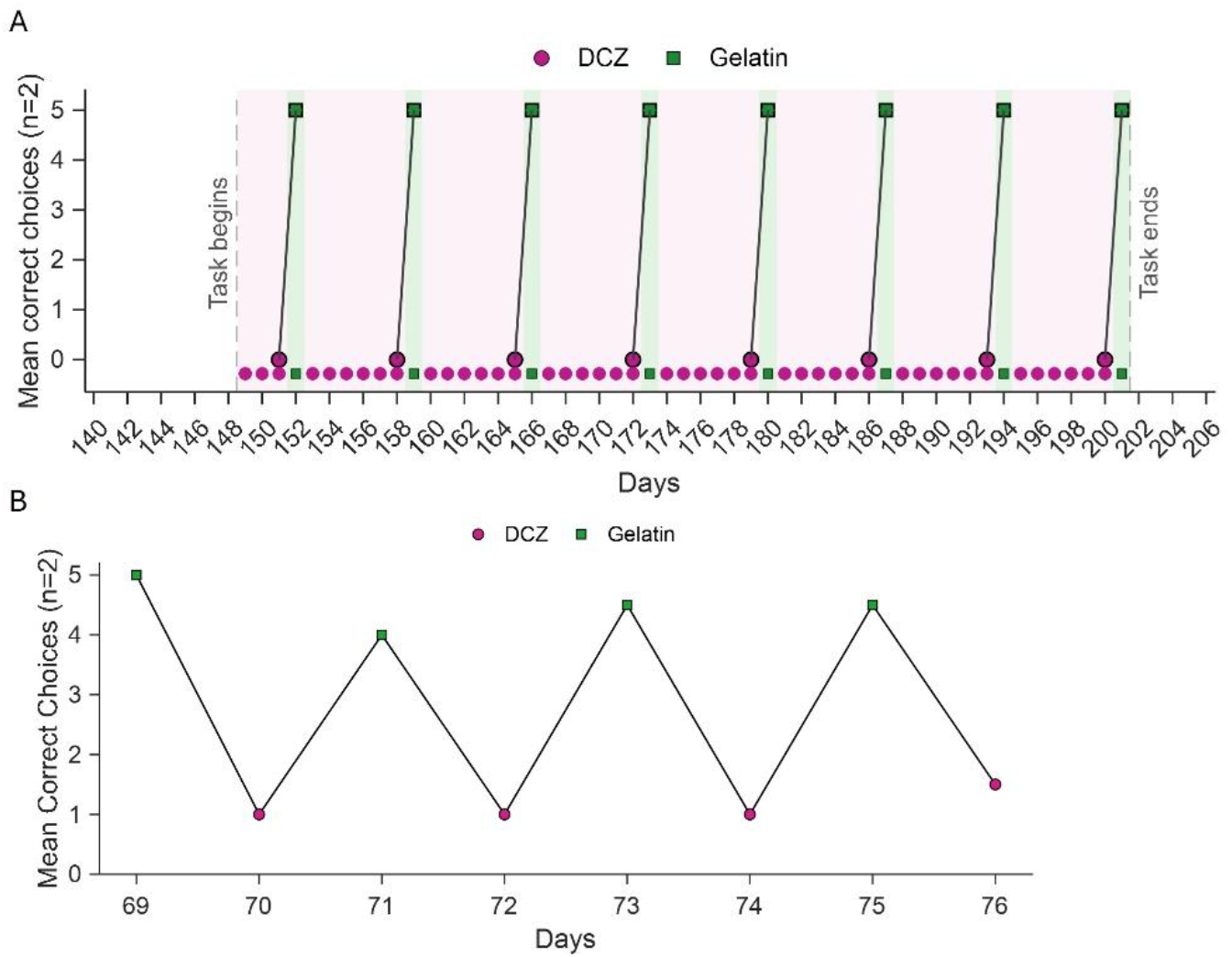
Behavioral efficacy of DCZ was maintained across 200 days of repeated administration and remained evident when behavioral testing began 1 hour after administration. (A) Two mice received 20 DCZ treatment sessions distributed across Days 1–147 (sessions not shown). Beginning on Day 148, DCZ was provided daily in the home cage, except on behavioral testing days, when mice received DCZ on one test day and gelatin on the other, according to the experimental schedule. (B) Mean number of correct choices across five trials during alternating DCZ (magenta circles) and gelatin (green squares) sessions. Behavioral testing began 1 hour after DCZ or gelatin administration. Lines connect consecutive testing sessions; *n* = 2 mice. Thus, the DCZ effect persisted long enough that waning drug effects cannot explain the gradual development of MS independence

**Supplementary Figure 8.**
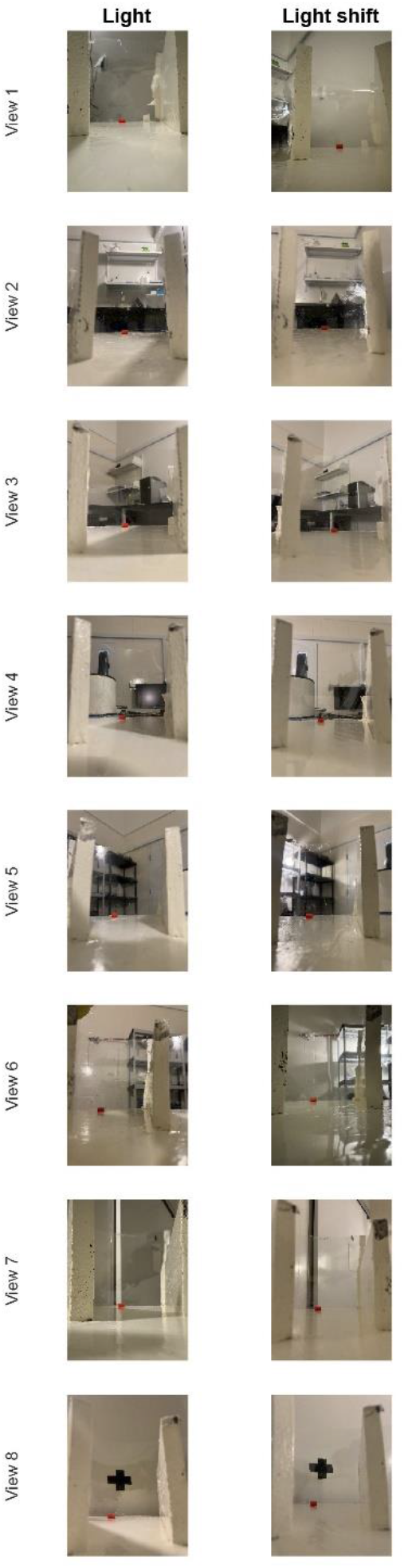
Visual environment before and after the 180° light shift. Representative views of the experimental room from each of the eight arms of the radial-arm maze. Left, views under the original light configuration. Right, corresponding views after the light was shifted by 180°.

